# Gene expression differences associated with intrinsic hindfoot muscle loss in the jerboa, *Jaculus jaculus*

**DOI:** 10.1101/2024.02.20.581295

**Authors:** Mai P. Tran, Daniel Ochoa Reyes, Alexander J. Weitzel, Aditya Saxena, Michael Hiller, Kimberly L. Cooper

**Affiliations:** Department of Cell and Developmental Biology, University of California San Diego, La Jolla, California 92093; LOEWE Centre for Translational Biodiversity Genomics, Senckenberganlage 25, 60325 Frankfurt, Germany; Senckenberg Research Institute, Senckenberganlage 25, 60325 Frankfurt, Germany; Goethe University Frankfurt, Faculty of Biosciences, Max-von-Laue-Str. 9, 60438 Frankfurt, Germany

## Abstract

Vertebrate animals that run or jump across sparsely vegetated habitats, such as horses and jerboas, have reduced the number of distal limb bones, and many have lost most or all distal limb muscle. We previously showed that nascent muscles are present in the jerboa hindfoot at birth and that these myofibers are rapidly and completely lost soon after by a process that shares features with pathological skeletal muscle atrophy. Here, we apply an intra- and inter-species approach, comparing jerboa and mouse muscles, to identify gene expression differences associated with the initiation and progression of jerboa hindfoot muscle loss. We show evidence for reduced Hepatocyte Growth Factor (HGF) and Fibroblast Growth Factor (FGF) signaling and an imbalance in nitric oxide signaling; all are pathways that are necessary for skeletal muscle development and regeneration. We also find evidence for phagosome formation, which hints at how myofibers may be removed by autophagy or by non-professional phagocytes without evidence for cell death or immune cell activation. Last, we show significant overlap between genes associated with jerboa hindfoot muscle loss and genes that are differentially expressed in a variety of human muscle pathologies and rodent models of muscle loss disorders. All together, these data provide molecular insight into the mechanism of evolutionary and developmental muscle loss in jerboa hindfeet.

## Introduction

Skeletal muscles produce force, pulling on the levers of bone to move the vertebrate body. Since locomotion is diverse across species (e.g., flying, running, jumping, swimming), so too are the sizes, shapes, and numbers of muscles that control bone movements. Many species that run or jump, such as large hooved mammals and saltatorial rodents, have substantially reduced the number of distal limb muscles that are no longer necessary for grasping and climbing. We previously showed that the three-toed jerboa (*Jaculus jaculus*), a desert adapted bipedal rodent, has lost all intrinsic muscles of the hindfoot over both evolutionary and developmental timescales (Tran et al., 2019). Although newborn jerboas have formed nascent myofibers of a single *m. flexor digitorum brevis* and three pinnate *m. interossei*, these myofibers begin to disappear by postnatal day 4 (P4) and are entirely absent in adults (Figure 1A, B).

**Figure 1:**
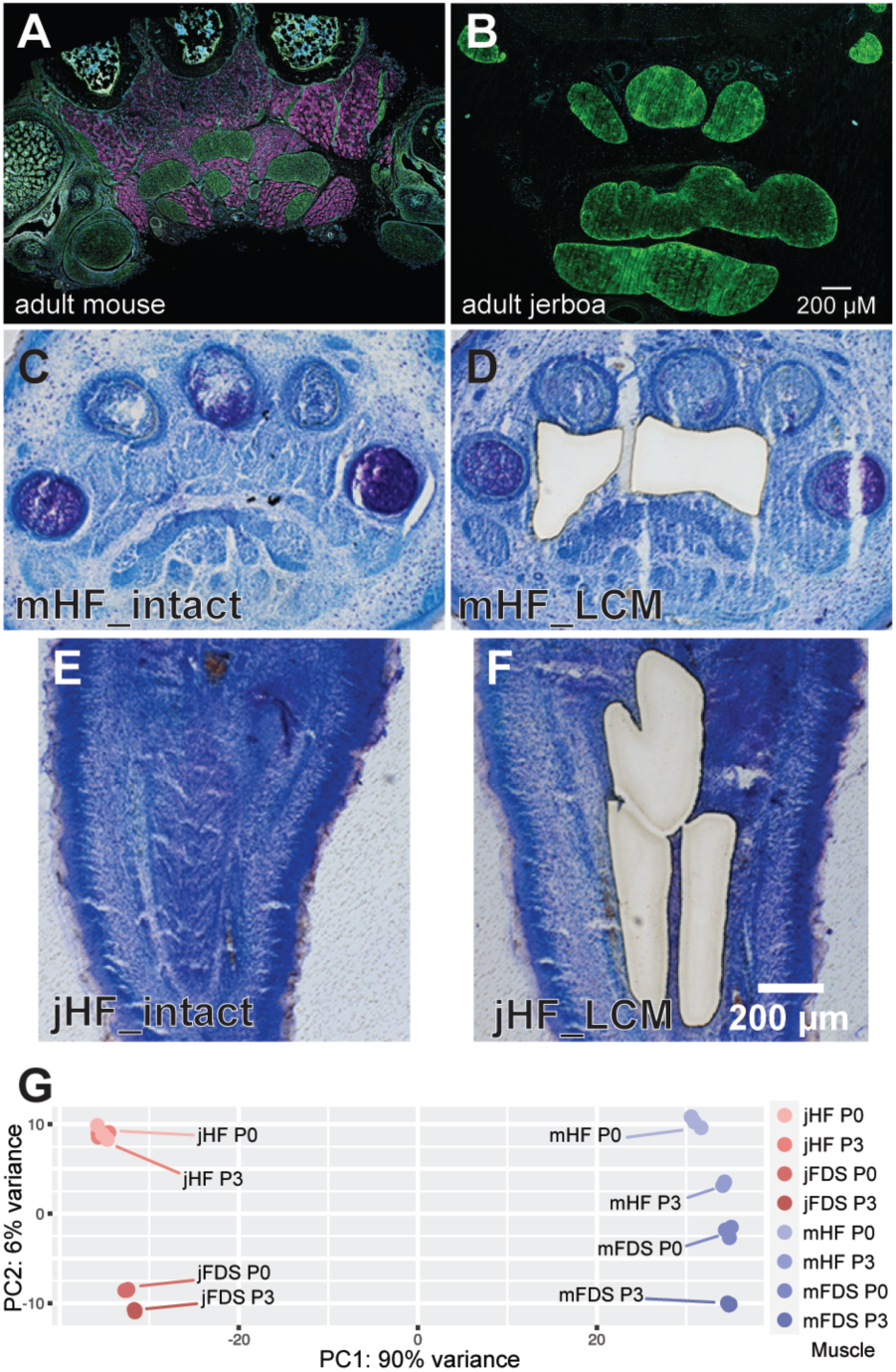
Sample morphology and preparation for differential mRNA expression analyses. (**A**, **B**) Transverse section through the mid-foot of adult mouse and jerboa hindfeet with immunofluorescent detection of pro-Collagen I (tendon, green) and skeletal muscle Myosin Heavy Chain (magenta). (**C**-**F**) Representative toluidine blue-stained plantar sections of mouse (mHF) or jerboa hindfoot (jHF) at P0 that are intact (**C**, **E**) or after laser capture microdissection (LCM) of intrinsic hindfoot muscle (**D, F**). (**G**) Principal components analysis of all jerboa and mouse hindfoot and flexor digitorum superficialis (FDS) transcriptomes.

Surprisingly, we found no evidence of apoptotic or necrotic cell death and no stimulation of a local immune response during stages of peak myofiber loss, countering well-supported models of developmental tissue remodeling (Tran et al., 2019). Instead, it appears that the immature contractile apparatus is disassembled in a stereotyped manner with Desmin being the earliest protein to become disorganized. The step-wise disassembly of the sarcomere, which is similar to its orderly disassembly during skeletal muscle atrophy, was associated with upregulation of E3 ubiquitin ligases that are also a hallmark of atrophy, *MuRF1* and *Atrogin-1*. However, skeletal muscle atrophy is typically considered a pathology associated with disuse, injury, starvation, or disease and typically causes a reduction in the size of individual myofibers but not their number (Moschella and Ontell, 1987).

Here, we implement an intersectional cross-species differential RNA-sequencing approach to broaden our understanding of molecular mechanisms that might be important for initiating and driving the unusual ‘atrophy-like’ process of muscle loss in the jerboa hindfoot. We use the laboratory mouse (*Mus musculus*) as a reference species; mice and jerboas diverged from a last common ancestor about 50 million years ago, and mice retained intrinsic hindfoot musculature typical of most other rodents. To account for gene expression divergence over such a long timescale that is likely unrelated to the mechanism of muscle loss in jerboas, we also sequenced RNA extracted from an analogous forelimb muscle that is retained in both species.

By intersecting gene expression differences within and between species at two timepoints, we identified sets of genes associated with the initiation and progression of jerboa hindfoot muscle loss. Among the significantly enriched genetic networks and pathways, we find evidence for lower Hepatocyte Growth Factor (HGF) and Fibroblast Growth Factor (FGF) signaling in jerboa hindfoot muscle than in other muscles that are retained. There is also evidence for an imbalance in the nitric oxide/arginine cycle suggesting lower nitric oxide signaling in jerboa hindfoot muscle. In addition to these pathways, which are known to be critical for muscle development, maintenance, and/or repair, we find evidence for phagosome formation suggesting a mechanism whereby remnants of nascent muscle may be removed either by autophagy or by non-professional phagocytic cells. Finally, we show significant overlap between our dataset and several human muscle degenerative disorders and rodent models of muscle disease lending further support to suggest that evolutionary muscle loss resembles a pathological state.

## Results

### Sample selection and experimental design

We showed previously that there was no significant difference in the number of myofibers located within the third interosseous muscle between postnatal day 0 (P0, birth) and P2 in either jerboa or mouse (Tran et al., 2019). However, whereas there is a significant increase over two-day intervals from P2 to P8 in mice, there is substantial variance between individual jerboas at P4 and a subsequent decrease until almost all myofibers are lost by P8. We further showed that the largely nascent and immature structure of the skeletal muscle sarcomere is most similar in the intrinsic hindfoot muscle of mouse and jerboa at P0, preceding degeneration in the jerboa. We therefore chose to isolate and sequence mRNA of the intrinsic hindfoot muscles at P0 and at P3 to capture molecular events at the initiation and during the process of degeneration but prior to tissue loss.

Unlike the larger muscles of the proximal limb, the intrinsic hindfoot muscles are very small making it extremely difficult to manually dissect tissue for transcriptome analysis. We therefore used laser capture microdissection (LCM) to isolate and enrich the intrinsic muscles from sections of P0 and P3 jerboa and mouse hindfeet (Figure 1C-F). To obtain sufficient material for sequencing and analyses, we pooled samples collected from the right and left hindfeet of six individuals for each of three biological replicates of each species and time point.

We then developed an experimental design to identify gene expression differences that might provide molecular evidence in support of a mechanism of muscle loss. As we showed previously for limb growth cartilages (Saxena et al., 2022), direct comparison of the homologous intrinsic hindfoot muscles of jerboa and mouse will identify the plethora of expression differences that accumulated since the two species diverged from their last common ancestor about 50 million years ago, most of which are likely unrelated to muscle loss in jerboas. Yet substantial expression diversity among different healthy skeletal muscles within an individual mouse or rat (Terry et al., 2018) suggests it would also be difficult to rely solely on direct comparison to a ‘typical developing’ jerboa muscle. Our approach therefore uses both within-species and between-species comparisons of jerboa hindfoot muscle that will be lost to muscles that will be retained in order to identify gene expression differences that are robustly associated with muscle loss.

We first sought an analogous forelimb muscle that is retained in both species. The intrinsic muscles of the hand are even smaller than in the hindfoot and thus more difficult to isolate in sufficient quantity. We therefore chose the *flexor digitorum superficialis* (FDS), which originates in the forelimb autopod during embryogenesis and later translocates to the fetal forearm (Huang et al., 2013). It is therefore evolutionarily and developmentally analogous to the intrinsic hindfoot muscles and also much larger and easy to manually dissect. We extracted mRNA from the FDS of one individual per three biological replicates of stage-matched (P0 and P3) jerboas and mice. We then processed all samples using the Illumina TruSeq Stranded mRNA Library Preparation Kit with polyA-enrichment and sample indexing and sequenced pools of indexed libraries using the Illumina HiSeq 4000 High Output platform.

### Differential expression analyses and filtering

For differential expression analyses that compare species transcriptomes directly, it is important to use a strongly supported 1:1 orthologous index of transcripts. We therefore applied TOGA, a method that uses a whole genome alignment to annotate coding genes, identifies (co)orthologous genes, and detects genes with reading frame inactivating mutations (Kirilenko et al., 2023). Using the *Mus musculus* genome (mm10) as reference and the revised *Jaculus jaculus* genome (mJacJac1.mat.Y.cur) as query, we annotated 16,667 1:1 orthologous transcripts in the two genomes from which we selected the longest isoform as representative of the gene body. We mapped sequenced reads from each biological replicate to the respective indexed genome. Principal component analysis (PCA) and sample-to-sample distance show segregation between experimental groups first by species and then by muscle type: jerboa hindfoot (jHF), jerboa FDS (jFDS), mouse hindfoot (mHF), mouse FDS (mFDS) (Figure 1G).

We then used DESeq2 to quantify differential expression between the hindfoot and FDS muscles of the jerboa at each stage (intra-species). We also quantified differential expression between jerboa and mouse hindfoot muscles and between jerboa and mouse FDS (inter-species) at each stage accounting for species specific transcript length normalization (Saxena et al., 2022). Statistically significant differentially expressed 1:1 orthologous transcripts in each pairwise analysis are defined as those with a p-adjusted (padj) value less than 0.05 (Supplementary Table 1). We did not apply a fold-change threshold, because genes with different functions (e.g., transcription factors versus enzymes) are likely differentially sensitive to altered expression levels.

To identify gene expression differences between species (inter-species) that are associated with jerboa hindfoot muscle loss, we first selected all genes that are significantly differentially expressed between jerboa and mouse in the hindfoot but not in the FDS (1,632 at P0; Figure 2A). We then plotted the log_2_ fold-change values for all genes that are significantly differentially expressed between species in both muscles (Figure 2B). The slope of the linear regression is 0.96 (R^2^=0.86), suggesting these genes have expression differences between species that are largely the same in both locations and likely unrelated to muscle loss specific to the jerboa hindfoot. However, 306 genes lie outside the 99% confidence interval; we therefore consider their expression differences between species to be ‘disproportionate’ in the two muscles. Combining the genes that are differentially expressed at P0 in hindfoot but not FDS with those that are disproportionately differentially expressed in hindfoot compared to FDS gives us 1,938 genes associated with muscle loss after the interspecies comparison. An identical filtering of samples collected from P3 muscles reveals 2,184 significantly differentially expressed genes are associated with muscle loss after the interspecies comparison at this later stage (Figure 3A, B).

**Figure 2:**
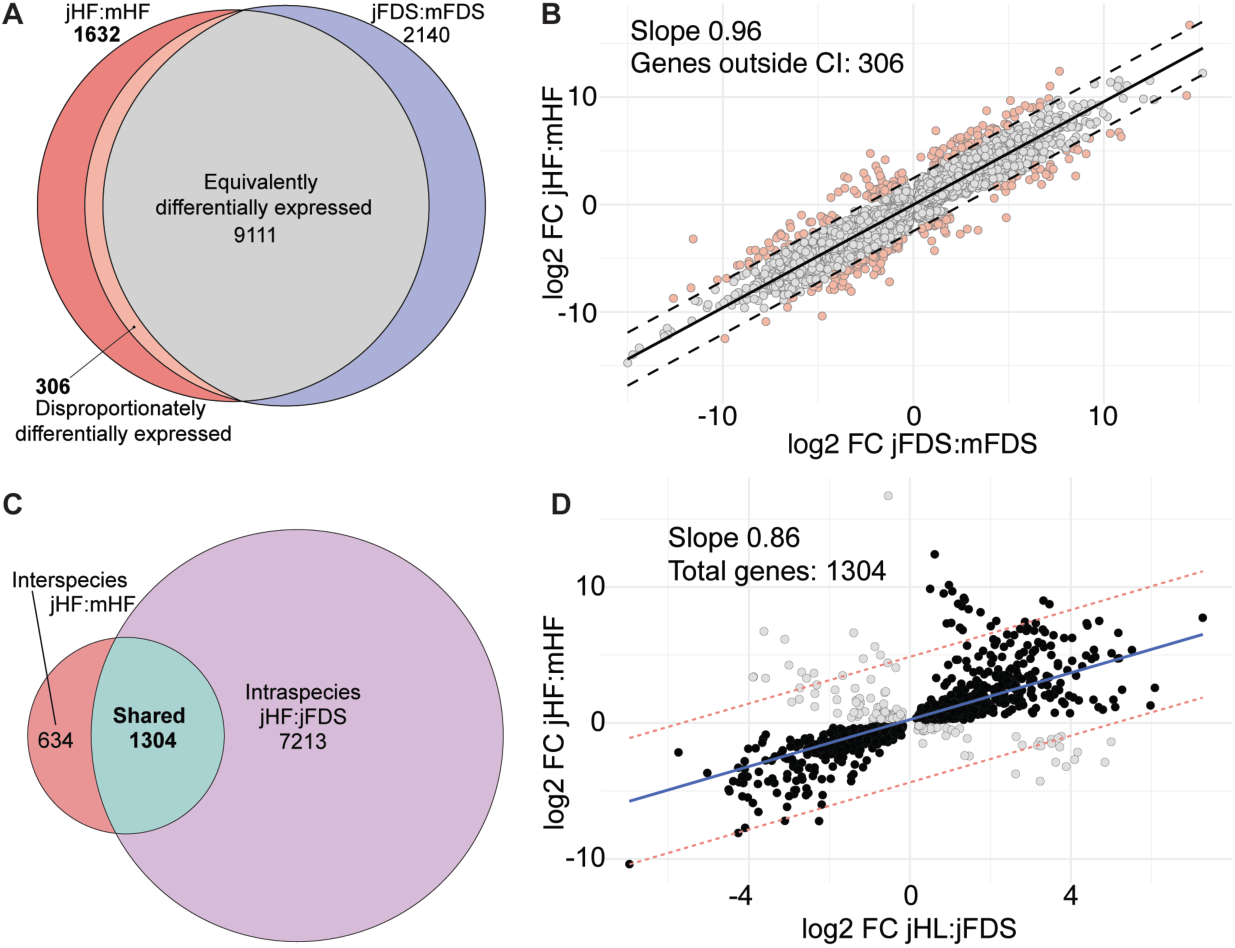
Identification of genes associated with the initiation of jerboa hindfoot muscle loss at P0. (**A**) Intersection of all genes that are differentially expressed between jerboa and mouse hindfoot muscle and between jerboa and mouse FDS. The orange sliver of ‘disproportionately differentially expressed’ genes lie outside of the 99% confidence interval of the regression of jerboa:mouse FDS versus hindfoot shown in (**B**). (**C**) The intersection of interspecies and intraspecies expression differences reveals genes that are differentially expressed in both comparisons. (**D**) A majority of differential expression correlates in the two comparisons; anti-correlated genes (gray dots) were removed.

**Figure 3:**
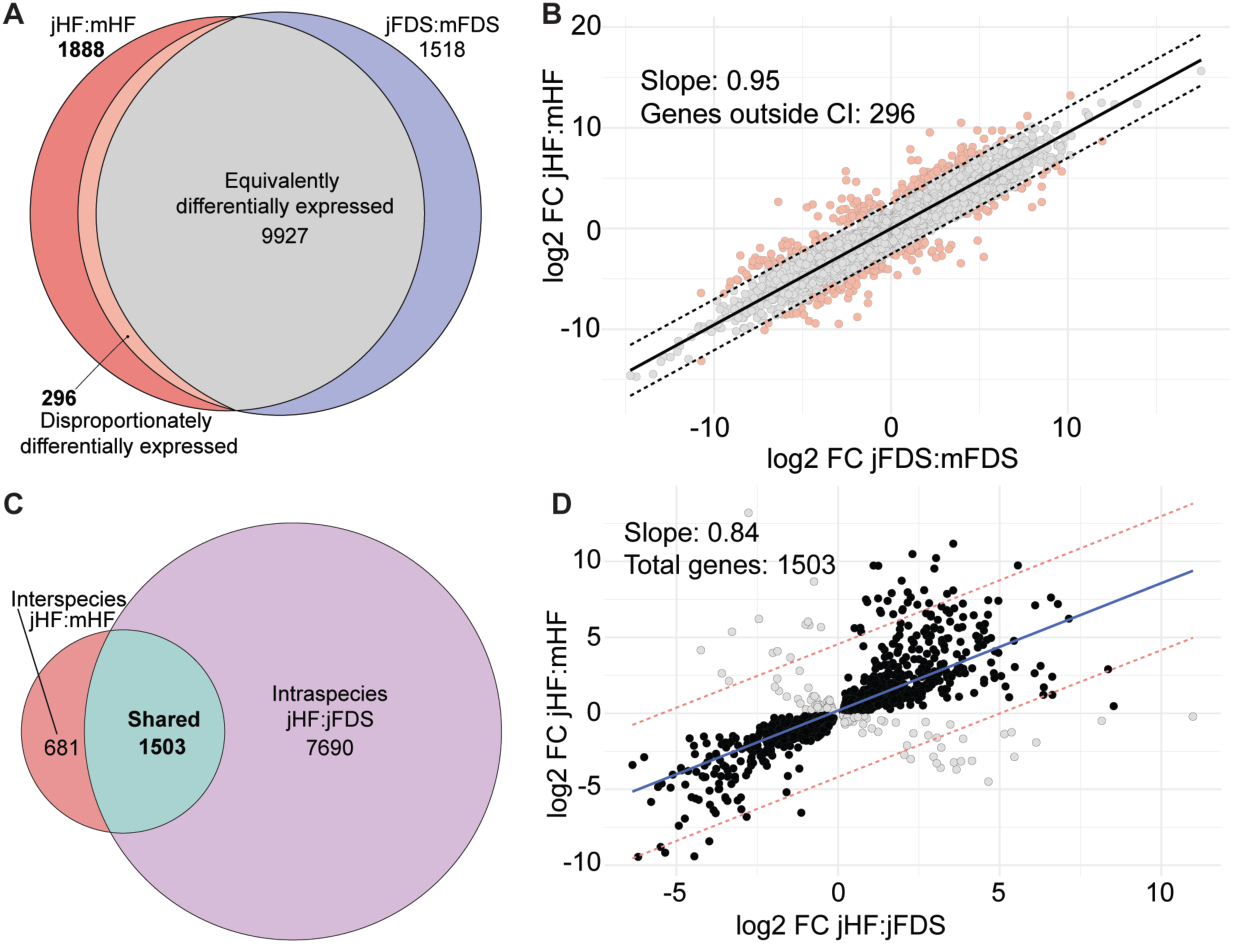
Identification of genes associated with the progression of jerboa hindfoot muscle loss at P3. (**A**) Intersection of all genes that are differentially expressed between jerboa and mouse hindfoot muscle and between jerboa and mouse FDS. The orange sliver of ‘disproportionately differentially expressed’ genes lie outside of the 99% confidence interval of the regression of jerboa:mouse FDS versus hindfoot shown in (**B**). (**C**) The intersection of interspecies and intraspecies expression differences reveals genes that are differentially expressed in both comparisons. (**D**) A majority of differential expression correlates in the two comparisons; anti-correlated genes (gray dots) were removed.

We next used the difference in developmental outcome of muscles within jerboas as a ‘second pass’ filter to identify genes that are also differentially expressed between jerboa hindfoot muscles that will be lost and FDS muscles that are retained (Figure 2C, 3C). We found correlations between the inter- and intraspecies expression differences with slope 0.86 (R^2^=0.42) at P0 and 0.84 (R^2^=0.48) at P3 (Figure 2D, 3D). We then selected all genes with consistent expression differences in the same direction in jerboa hindfoot muscle that is lost compared to mouse hindfoot and jerboa FDS muscles that are retained. Altogether, these inter- and intraspecies analyses identified 1162 genes associated with jerboa hindfoot muscle loss at P0 and 1382 genes at P3 (black dots in Figure 2D and 3D; Supplementary Table 2), which we used for all subsequent candidate gene and network and pathway analyses. Comparing the two timepoints, we find that 749 genes are differentially expressed only at P0, 969 are differentially expressed only at P3, and 413 genes are differentially expressed at both timepoints. Among these that are consistent, all but two differ in the same fold-change direction at both stages (Supplementary Figure 1).

### Mechanistic insights from gene expression differences and pathway enrichment analyses

These gene sets provide an opportunity to explore possible mechanisms of evolutionary and developmental muscle loss in the jerboa hindfoot. We first implemented a network and pathway enrichment analysis of all genes associated with jerboa hindfoot muscle loss at P0 and at P3 using Ingenuity Pathway Analysis (IPA, Qiagen). Canonical pathway analysis of well-characterized metabolic and cell signaling pathways in IPA showed significant enrichment [-log(p-value) >1.3] for 118 pathways at P0 and 32 pathways at P3. The 20 most significantly enriched pathways at each time point are presented in Figure 4, and all significant pathways are in Supplementary Table 3. Here, we focus on a few notable differentially expressed genes and pathways that functionally relate to muscle development, regeneration, and/or maintenance, providing insight into the possible molecular mechanisms of jerboa hindfoot muscle loss.

**Figure 4:**
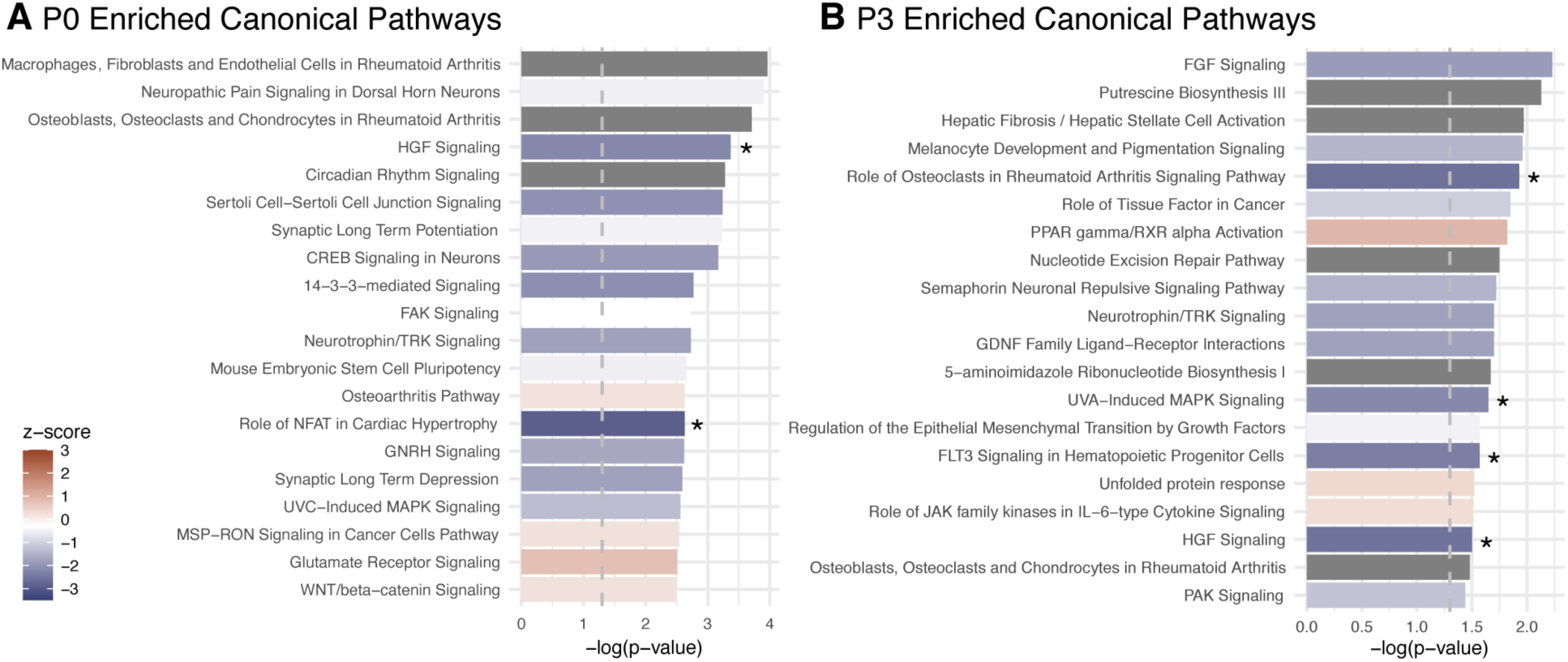
The top twenty most significantly enriched canonical pathways among genes associated with jerboa hindfoot muscle loss at P0. (**A**) **and at P3** (**B**). Vertical dashed lines mark the threshold for significance [-log(0.05)=1.3]. Asterisks mark pathways that reach significance for ‘activation’ (z-score >2) or inhibition (z-score<-2) reported within the Ingenuity Pathway Analysis.

We previously observed no evidence of cell death by a variety of markers and no macrophages in the vicinity of jerboa hindfoot muscles during degeneration (Tran et al., 2019). It is therefore unclear how nascent myofibers disappear after showing signs of ‘atrophy-like’ degeneration. Here, we show that IPA calls the ‘Phagosome Formation’ canonical pathway as significantly enriched at both P0 and P3, and the ‘Unfolded Protein Response’ pathway as enriched at P3. Absence of evidence for professional phagocytic cells (e.g., macrophages and dendritic cells) in our previous work suggests that phagosomes might form in another cell type. It is possible the enriched phagosome formation pathway reflects myofiber autophagy, whereby muscle cells may degrade and recycle their own damaged proteins (Xia et al., 2021), which could be consistent with the Unfolded Protein Response. Alternatively, phagosomes may form within fibroblasts that we previously observed intermingled with highly degenerating muscle by electron microscopy and immunofluorescence (Tran et al., 2019). If so, this would suggest these are non-professional phagocytic cells that might consume the remains of myofibers.

The ‘HGF Signaling’ pathway appears to be significantly inhibited (z-score <-2) in jerboa hindfoot muscle at both P0 and P3 based on differential expression of networked molecules. This result stands out as notable due to the well-established importance of the HGF ligand and c-Met receptor to multiple aspects of muscle cell biology. Homozygous *c-met* loss-of-function mice lack all limb muscle, as well as muscles of the diaphragm and tip of the tongue, due to defective muscle precursor migration (Bladt et al., 1995). In embryonic chickens, exogenous HGF is sufficient to stimulate muscle precursor migration and also prevents myogenic differentiation (Scaal et al., 1999). The importance of HGF/c-Met signaling is not limited to embryogenesis; HGF is released by injured adult muscle and stimulates the c-Met receptor expressed by satellite cells (Tatsumi et al., 1998; Miller et al., 2000). Satellite cells are quiescent muscle stem cells nestled between the muscle and basal lamina, which are activated by HGF signaling to re-enter the cell cycle and become migratory. These cells then fuse to one another to form new myofibers or to injured myofibers for repair. Thus, HGF signaling is also essential to the earliest stages of muscle regeneration after injury. Furthermore, evidence suggests that HGF can inhibit or reverse skeletal muscle atrophy induced by denervation and that cMet inhibition after nerve injury further increases expression of the E3 ubiquitin ligases, *Murf1* and *Atrogin1* (Choi et al., 2018). Altogether, these findings suggest that evidence for HGF pathway inhibition is consistent with a putative role in the rapid loss of jerboa hindfoot muscle.

‘FGF Signaling’ is the most significantly enriched pathway at P3 with a z-score trending toward significant inhibition. At least six ligands and two receptors of this highly pleiotropic growth factor pathway are expressed in the skeletal muscle lineage of mouse and/or rat (Hannon et al., 1996; Kästner et al., 2000), though most loss-of-function mice have normal or minimally affected skeletal muscle possibly due to redundancies (Pawlikowski et al., 2017). An exception, *Fgf6*, appears to be necessary for an early postnatal expansion of the muscle stem cell pool, which may affect muscle regeneration after injury (Floss et al., 1997; Zofkie et al., 2021). Notably, *Fgf6* ligand expression in jerboa hindfoot muscle is 4.7-fold lower than in mouse hindfoot and 6.8-fold lower than in jerboa FDS. Although *Fgf6* is also significantly differentially expressed in jerboa hindfoot at P0 (3.3-fold and 4.1-fold lower than mouse hindfoot and jerboa FDS, respectively), the FGF signaling pathway is not significantly enriched at this earlier stage.

Considering also the largest fold-change differences in each dataset reveals that *Nitric oxide synthase 1* (*Nos1/nNos*) is expressed a hundred to a thousand-fold lower at P0 in jerboa hindfoot muscle when compared to either mouse hindfoot muscle (log_2_ fold-change = −10.4; padj = 1.7E-17) or jerboa FDS (log_2_ fold-change = −7.0; padj = 2.7E-07) (Figure 2; Supplementary Table 2). IPA calls the ‘nNos Signaling in Skeletal Muscle Cells’ canonical pathway as significantly enriched at P0 (-log_10_ p-value=1.33). Nitric oxide (NO) is a gaseous molecule with important functions in many tissues (Lundberg and Weitzberg, 2022). In skeletal muscle, NO regulates key aspects of cell biology and physiology, including early stages of myogenesis, muscle force production, metabolism, and repair after muscle injury (Stamler and Meissner, 2001). Nitric oxide is produced by the catalytic activity of *Nos1*, which converts L-Arginine to NO and L-Citrulline. As evidence of the importance of NO signaling in muscle maintenance and repair, *Nos1* activity is reduced in multiple muscle degenerative disorders with a variety of genetic underpinnings (Brenman et al., 1995; Chao et al., 1996; Crosbie et al., 2002). Furthermore, *Nos1*^-/-^ knockout mice have a smaller myofiber cross-sectional area, reduced force production, and show ultrastructural damage to the sarcomere after exercise (De Palma et al., 2014). Interestingly, *Argininosuccinate synthase 1* (*Ass1*) has one of the largest fold-change differences among genes that are expressed higher in jerboa hindfoot muscle at both P0 (log_2_ fold-change >4.7) and at P3 (log_2_ fold-change >5.6) when compared to muscles that are retained in each species. *Ass1* catalyzes a key step in the biosynthesis of cellular L-Arginine from L-Citrulline, the secondary product of *Nos1* activity (Wu and Morris, 1998). Together, this suggests that an imbalance in the nitric oxide/arginine cycle might also contribute to jerboa hindfoot muscle loss.

### Comparison of genes associated with evolutionary muscle loss and models of pathological muscle loss

Pathological muscle loss can result from disease causing mutations or in response to denervation, disuse, cancer, fasting, or aging. Are the molecular mechanisms of muscle loss in the jerboa hindfoot broadly similar to pathological muscle loss or similar to a narrower subset of disorders? To answer this question, we selected twenty-four publicly accessible differential expression (RNA-Seq or microarray) datasets that compared human biopsies or mouse or rat models of pathological muscle loss to healthy control skeletal muscle. Datasets were included only if the full list of differentially expressed genes reported were accessible without requiring reanalysis of the raw data.

We first identified differentially expressed genes (p-adj<0.05) within each of the mouse, rat, and human datasets that were assigned the same name (and unique ENMUSG for mouse genes) as in our 1:1 jerboa/mouse orthologous reference set. Three datasets were excluded at this point because fewer than 50 differentially expressed genes remained after filtering for jerboa/mouse orthologs. Using the 16,667 jerboa and mouse 1:1 orthologs as the total number of genes, we performed a Fisher’s exact test with Benjamini-Hochberg multiple hypothesis correction to identify significant overlap between each disease/pathology dataset and the sets of genes associated with jerboa hindfoot muscle loss at P0 and at P3 (Table 1). Of the 21 pathology datasets, we found that four overlap significantly with jerboa hindfoot muscle loss at P0, five overlap significantly with jerboa hindfoot muscle loss at P3, and fourteen do not significantly overlap with jerboa hindfoot muscle loss at either developmental stage.

**Table 1:**
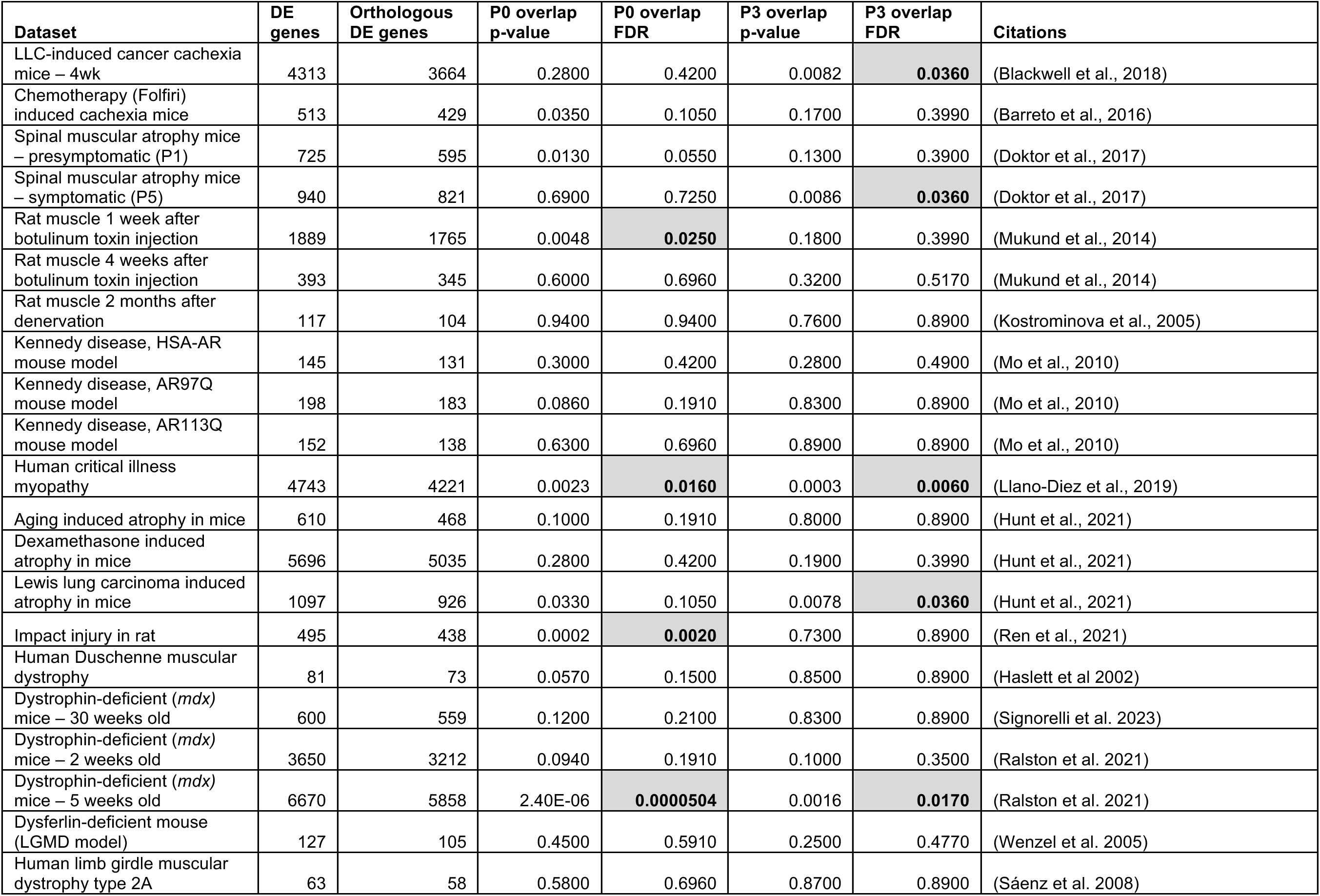
Values for the Fisher’s Exact Test of overlap between genes associated with jerboa hindfoot muscle loss at P0 or at P3 and datasets obtained from rodent disease models or human pathologies. Significant values are bold and highlighted.

Two pathology models overlap significantly at both timepoints: human critical illness myopathy (CIM) and the mdx mouse model of Duchenne’s muscular dystrophy (Llano-Diez et al., 2019; Ralston et al., 2021). Critical illness myopathy (CIM), also known as acute quadriplegic myopathy, is the significant depletion of skeletal muscle mass and compromised performance in individuals receiving intensive care (Latronico et al., 1996; De Jonghe et al., 2002). The underlying mechanisms of CIM are not fully understood but involve processes such as activation of protein degradation pathways, decreased expression of myofibrillar proteins, reduced excitability of cell membranes, mitochondrial dysfunction, and altered excitation-contraction coupling (Shepherd et al., 2017). Duchenne’s muscular dystrophy (DMD), on the other hand, is one of the most well-characterized and severe forms of hereditary muscular dystrophy. DMD is caused by mutations in *Dystrophin*, a large protein component of the complex linking and stabilizing the myofiber cytoskeleton to the extracellular matrix. The *mdx* mouse model has a spontaneous mutation that prematurely terminates *Dystrophin* translation (Bulfield et al., 1984; Ryder-Cook et al., 1988; Sicinski et al., 1989), and it is one of the most widely used rodent models of human DMD.

At P0 but not P3, we see overlap with a botulinum toxin rat model of atrophy one week after treatment, and with a rat skeletal muscle injury model (Mukund et al., 2014; Ren et al., 2021). Injection of BT was used to inhibit motor neuron activity, thus mimicking conditions of muscle inactivity often seen in multiple neuromuscular disorders or bed-ridden patients (Mukund et al., 2014). The authors performed a long-term study of BT-induced muscle loss from one week to up to a year after BT injection and reported that the most dramatic transcriptome changes (1989 genes) occurred within one week compared to four weeks or longer. The transcriptional differences reported after mechanical injury to the rat tibialis anterior were observed within hours of wounding (Ren et al., 2021).

At P3 but not P0, we see significant overlap between genes associated with jerboa hindfoot muscle loss and differentially expressed genes in two independent mouse models of cancer cachexia (Blackwell et al., 2018; Hunt et al., 2021) and a symptomatic model of spinal muscular atrophy (Doktor et al., 2017). Cancer-induced cachexia is a highly complex metabolic syndrome characterized by progressive muscle wasting (Fearon et al., 2012). Notable clinical manifestations of cachexia include loss of weight, inflammation, resistance to insulin, and heightened breakdown of muscle proteins (Fearon et al., 2012). In both mouse models, cancer-induced muscular atrophy was caused by injection of Lewis lung carcinoma (LLC) mouse tumor cells, which lead to the reduction in size of type 2B myofibers with no change in the number of myofibers or the relative distribution of different myofiber types (Hunt et al., 2021).

Spinal muscular atrophy is a neuromuscular disease caused by deficiency of the ‘Survival of Motor Neurons’ (SMN) protein, which leads to progressive muscle weakness and often causes death in infancy. In a mouse model of severe SMA, animals have a rapid disease progression and a median lifespan of 10 days. Transcriptome analyses at P1 (pre-symptomatic) and P5 (symptomatic) identified hundreds of differentially expressed genes in SMA skeletal muscle compared to control (Doktor et al., 2017). We find significant overlap between our P3 jerboa hindfoot muscle loss dataset and the symptomatic P5 SMA mouse dataset. Together, these intersections lend support to a hypothesis that jerboa hindfoot muscle loss progresses with a gene expression profile similar to pathological atrophy. That only a subset of pathologies overlap with jerboa hindfoot muscle loss is consistent with observations that different causes of skeletal muscle atrophy also have minimal overlap with one another (Hunt et al., 2021).

### Conclusions and Limitations

Individual muscles and groups of muscles have been lost repeatedly throughout vertebrate phylogeny as evolution has reshaped the musculoskeletal system to enable a variety of types of locomotion. We previously reported that the cellular mechanisms of intrinsic hindfoot muscle loss in neonatal jerboas, which is histologically similar to intrinsic hindfoot muscle loss in fetal horses (Cunningham, 1883), have atrophy-like characteristics. Here we applied a comparative transcriptomics approach comparing gene expression in jerboa hindfoot muscles to both mouse hindfoot muscles (inter-species) and to jerboa forelimb muscle (intra-species), which are retained to adulthood. Genes we identified with consistent expression differences (same fold-change direction) in these multi-way comparisons are therefore ‘associated with jerboa hindfoot muscle loss’. The complete datasets are available in Supplementary Tables 1 and 2.

Correlation of gene expression differences between inter- and intraspecies analyses and between stages of muscle loss (Supplementary Figure 1) supports the logic of our experimental design, but we acknowledge that the datasets are likely incomplete and include false positives. Furthermore, these are snapshots of gene expression differences that include genes that may be causative and others that are certainly a consequence of the primary mechanism of muscle loss due to the interconnected effects of expression perturbation (Cowen et al., 2017). Nevertheless, these datasets provide evidence for molecular mechanisms of evolutionary loss of distal limb skeletal muscles.

Although application of pathway enrichment analyses to these datasets is limited by the current knowledge base of gene functions, it does provide valuable insight into the putative molecular mechanisms of a biological phenomenon. Here, we have drawn attention to a few enriched pathways (HGF, FGF, and NO) with well-documented importance to muscle development, maintenance, and/or repair; all enriched pathways are available in Supplementary Table 3. As with all such ‘omics’ datasets, it will be important to repeat these analyses as the functional annotation of all genes continues to expand, thus enabling broader and deeper insight into putative molecular mechanisms of a variety of biological processes.

Finally, we set out to determine if these differential expression analyses would provide further support for a hypothesis that the mechanism of jerboa hindfoot muscle loss, occurring over both evolutionary and developmental timescales, shares substantial similarities with instances of pathological muscle loss. At one or both stages of jerboa hindfoot muscle loss, we found significant overlap with seven out of 21 of the analyzed disease and injury states, consistent with our prior histological and ultrastructural observations. However, we emphasize that statistically significant overlap is evidence for *correlation* of the gene expression differences observed in jerboa hindfoot muscle and certain muscle pathologies, but this should not be interpreted to suggest that the *causes* are the same. Rather, these correlations with specific pathology states demonstrate striking and perhaps surprising similarity between evolutionary and pathological states while also serving as entry points to gain further insight into the mechanisms of muscle loss in a variety of contexts.

## Materials and Methods

### Animals

Jerboas were housed and reared as previously described (Jordan et al., 2011). Outbred CD1 mice were obtained from Charles River Laboratories (MA, USA), housed in standard conditions, and fed a breeder’s diet. All animal care and use protocols for mice and jerboas were approved by the Institutional Animal Care and Use Committee (IACUC) of the University of California, San Diego.

### Laser capture microdissection of hindfoot muscles and bulk dissection of FDS

Mouse and jerboa feet were fresh frozen in blocks of OCT freezing media, and blocks were stored at −80**°**C until cryosectioned. Blocks were sectioned at 30 µm thickness, and sections were transferred to PEN (polyethylene naphthalate) MembraneSlides (Zeiss). Sections were stained and dehydrated using the Arcturus HistoGene LCM Frozen Section Staining Kit following manufacturer’s protocol (ThermoFisher). Sections were subjected to laser microdissection using Zeiss PALM MicroBeam microscope according to manufacturer’s instructions. Immediately after capture, we added extraction buffer from the PicoPure RNA Isolation kit and incubated samples for 30 minutes at 42**°**C prior to storage at −80**°**C until RNA isolation. Tissues were pooled from six animals (right and left limbs) per biological replicate and three biological replicates were prepared per species and stage (mouse and jerboa, P0 and P3). To isolate the FDS muscle in each species, we removed and skinned each forelimb, located the tendons of the *m. flexor digitorum superficialis*, and followed the tendon to the muscle in the forearm. We then severed the muscle at each tendon. Each biological replicate for the FDS is one animal (right and left FDS). Samples were incubated overnight in RNAlater (Invitrogen) prior to storage at −80**°**C until RNA isolation.

### mRNA isolation and sequencing

mRNA extraction was performed using the PicoPure RNA Isolation Kit (Thermo Fisher) according to the manufacturer’s instructions. RNA quality and concentration were determined using Agilent TapeStation (Agilent Technologies, Santa Clara, CA). RIN^e^ scores from TapeStation analysis for all samples used were at minimum 7.0. Libraries were prepared using the Illumina TruSeq Stranded mRNA Library Preparation Kit using a polyA-enrichment strategy and sample indexing. Two pools of mouse and jerboa samples were loaded onto each lane of Illumina HiSeq 4000 High Output flow cell and sequenced in a 1 × 75 bp single read format. RNA sequencing was completed at the Institute for Genomic Medicine core facility at UC San Diego (La Jolla, CA).

### RNA-Seq read mapping and differential expression

Adaptors and low quality bases were trimmed from sequences by using Trimmomatic with default parameters (Bolger et al., 2014). Quality control of sequences in FASTQ and BAM format was assessed with the FastQC software (Babraham Bioinfomatics, http://www.bioinformatics.babraham.ac.uk/projects/fastqc/). We then used the STAR aligner to map reads to the respective genome (mouse mm10 or jerboa mJacJac1.mat.Y.cur) (Dobin et al., 2013). Each genome was annotated using a 1:1 jerboa to mouse orthologous gene annotation set generated using TOGA (Kirilenko et al., 2023). Trimmomatic, FastQC, and STAR analysis were performed on Amazon Web Services EC2. Read counts associated with each specific transcript were used to carry out analysis of differential expression with DESeq2 (Love et al., 2014) with an additional transcript length normalization for each species in the interspecies comparison of homologous muscle (Saxena et al., 2022).

### Data Intersections and Filtration

The output of DESeq2 resulted in eight individual files: (jHF:mHF), (jFDS:mFDS), (jHF:jFDS), (mHF:mFDS) at each of P0 and P3. These initially contained expression values for 17,641 genes (all 1:1 orthologues plus genes designated as present in single copy in mouse and jerboa genomes but predicted non-functional in jerboa – one to zero). To these, we joined a column containing the gene name from a metadata .tsv file linking ENMUST to gene name. This .tsv file lacked the identities of 3 genes, *Tusc3*, *Kmt5b*, and *Wdfy1* that were added manually. The gene sets were then subset to contain only the 16,667 1:1 orthologous genes. Next, the 1:1 gene sets were subset to contain only genes with a p-adjusted value less than 0.05, resulting in:

- jFDS:mFDS P0 = 11557 differentially expressed 1:1 orthologs
- jFDS:mFDS P3 = 11741 “ “
- jHF:mHF P0 = 11049 “ “
- jHF:mHF P3 = 12111 “ “
- jHF:jFDS P0 = 8517 “ “
- jHF:jFDS P3 = 9193 “ “
- mHF:mFDS P0 = 7496 “ “
- mHF:mFDS P3 = 8359 “ “

Each of the following steps was replicated identically for data collected at the P0 and P3 timepoints. For interspecies comparisons, the gene set for jHF:mHF was intersected with jFDS:mFDS in order to obtain a list of genes that are significantly differentially expressed only between the hindfoot muscle of the two species (1632 at P0, 1888 at P3), as well as a set of genes that are differentially expressed between the hindfoot muscle and also between the FDS muscle of the two species (9417 at P0, 10223 at P3). After plotting log_2_ fold-change differential expression values for jFDS:mFDS versus jHF:mHF, we determined the 99% confidence interval to identify disproportionately differentially expressed genes that lie outside of this interval (306 at P0, 296 at P3). These disproportionately differentially expressed genes were added to the ‘hindfoot only’ differentially expressed genes to create the interspecies differential expression dataset (1938 genes at P0, 2184 genes at P3).

Next, the interspecies datasets were each intersected with the same stage intraspecies (jHF:jFDS) dataset to find genes commonly differentially expressed when jerboa hindfoot muscle is compared to both muscles that are retained. This intersection resulted in 1,304 genes at P0 and 1,503 genes at P3. Genes with expression differences that were anti-correlated (e.g., positive log_2_ fold-change in the interspecies comparison but negative log_2_ fold-change in the intraspecies comparison) were removed, resulting in final datasets of significant differentially expressed genes associated with jerboa hindfoot muscle loss (1,162 genes at P0 and 1,382 genes at P3). All graphs, data filtering, and statistical analyses reported in this manuscript were generated using the R-stats package.

### Ingenuity Pathway Analysis (Qiagen)

The reference file for use in IPA included only genes with a read count value greater than 1 in at least one of the 24 replicate samples: 3 jFDS, 3 mFDS, 3 jHF, 3 mHF at P0 and P3. This reference file contained 16,264 of the 16,667 1:1 orthologous genes. Graphs were made for canonical pathways from IPA generated data (Supplementary Table 3).

### Significance of overlap with other muscle atrophy models

We took each gene list from the original cited authors’ reported DESeq2 or microarray analysis output. We then selected genes with a differential expression adjusted p-value (DESeq2) or p-value (microarray) less than 0.05. Gene lists were then converted to ENSMUSG using g:Profiler gene ID Conversion function (https://biit.cs.ut.ee/gprofiler/convert). Genes with no associated ENSMUSG or an ENSMUSG not found in our jerboa to mouse 1:1 orthology dataset were removed from consideration. We performed all Fisher’s exact tests using the GeneOverlap package in R to determine statistical significance of overlapping genes (Shen). We used the jerboa to mouse 1:1 orthology dataset (16,667 genes) as the total gene set for Fisher’s exact tests. Adjusted p-values for multiple comparisons were calculated in R using the Benjamini & Hochberg method.

## Data Accessibility

The data discussed in this publication have been deposited in NCBI’s Gene Expression Omnibus (Tran *et al*., 2023) and are accessible through GEO Series accession number GSE235932 (https://www.ncbi.nlm.nih.gov/geo/query/acc.cgi?acc=GSE235932). Supplementary Excel tables are available at <u>10.5281/zenodo.10685550</u>

## Acknowledgements

We thank Dr. Uri Manor and Dr. Tong Zhang in the Biophotonics Core at the Salk Institute for Biological Studies for assistance and usage of the Zeiss PALM MicroBeam microscope. Illumina HiSeq library sequencing was completed by the Institute for Genomic Medicine Genomics Center at UC San Diego, and the Center for Computational Biology at UC San Diego provided bioinformatic assistance. We would also like to thank Dr. Kavitha Mukund for advice on bioinformatic tools and Dr Ronghui Xu and Dr. Lin Liu for statistical advice. This work was supported by the National Institute for Arthritis and Musculoskeletal and Skin Diseases under award number AR074609 and by the Wu Tsai Human Performance Alliance and the Joe and Clara Tsai Foundation. MPT was supported by the Cellular and Molecular Genetics Training Grant at UC San Diego, funded by the National Institute for General Medical Sciences under award number T32GM724039.

---

**Supplementary Figure 1:**
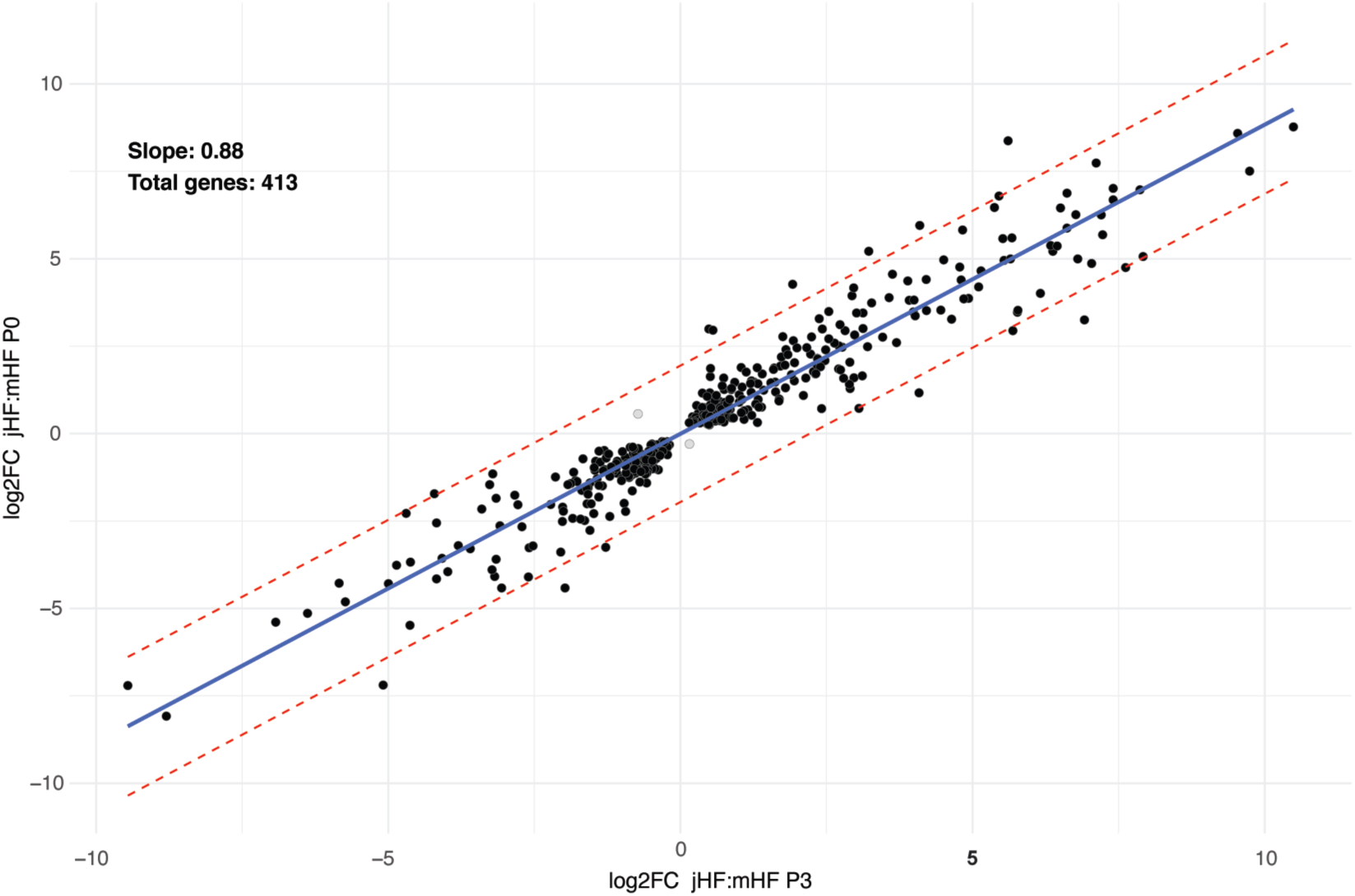
Strong correlation of gene expression differences between jerboa and mouse hindfoot muscle at PO and at P3. Only two genes (gray dots) are anti-correlated.

